# An error correction strategy for image reconstruction by DNA sequencing microscopy

**DOI:** 10.1101/2023.07.10.548317

**Authors:** Alexander Kloosterman, Igor Baars, Björn Högberg

## Abstract

By pairing adjacent molecules in-situ and then mapping these pairs, DNA microscopy could significantly reduce workload in spatial omics methods by directly inferring geometry from sequencing data alone. However, experimental artefacts can lead to errors in the adjacency data which distort the spatial reconstruction. Here, we describe a method to correct two such errors: spurious crosslinks formed between any two nodes, and fused nodes that are formed out of multiple molecules. We build on the principle that spatially close molecules should be connected and show that these errors violate this principle, allowing for their detection and correction. Our method corrects errors in simulated data, even in the presence of up to 20% errors, and proves to be more efficient at removing errors from experimental data than a read count filter. Integrating this method in DNA microscopy will significantly improve the accuracy of spatial reconstructions with lower data loss.

## Introduction

With the improvement in sequencing technology, techniques to investigate biological samples have become increasingly refined, progressing from sequencing in bulk, to single-cell RNA ^1^, to spatial transcriptomics ^2, 3^. The latter technique allows one to obtain the organization cells in tissue, leading to deeper insights in biology and improving the detection of diseases^4, 5^. However, current techniques for spatial transcriptomics rely on fluorescence microscopy, which is limited in throughput, especially for large amounts of targets^3^.

DNA microscopy is an emerging spatial transcriptomic technique that aims to find the spatial organization of DNA or RNA using sequencing alone, bypassing the use of optical microscopy. The common theme in all DNA microscopy methods is to use PCR and sequencing to find pairs of adjacent molecules use that to find their relative locations^6–9^ (Fig. 1). In a typical workflow (Fig. 1A), molecules are barcoded and amplified locally in tissue of interest, creating polymerase colonies (polonies)^10^ (Fig. 1B). Where polonies overlap, amplicons are engineered to fuse, forming concatemers (Fig. 1C) which, when sequenced, reveal which two polonies are adjacent. All the adjacency data is represented in a graph, where each node represents one polony, and each edge a sequenced concatemer. From this graph the original locations can be estimated^6, 7, 9^ (Fig. 1D). Importantly, the information of these adjacency pairs is obtained with sequencing information only, meaning that a) it can capture both the sequence and location of transcripts simultaneously and many targets can be captured simultaneously, b) it does not require processing or stitching of image data, only analysis of sequencing data and c) it is not inherently limited to 2D reconstructions but can be used to reconstruct 3D samples as well.

**Figure 1.**
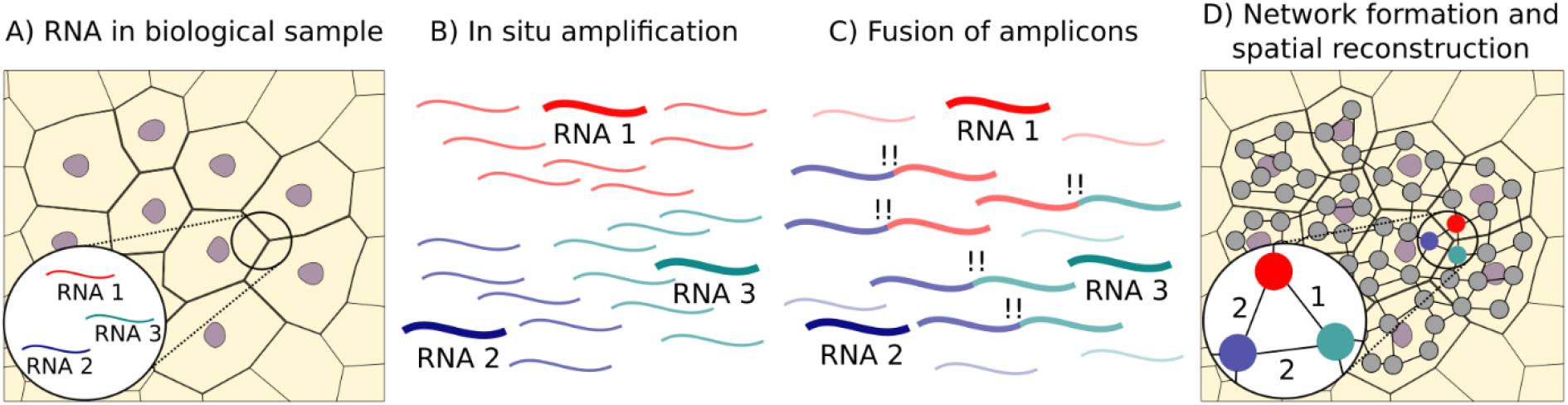
DNA microscopy reveals spatial locations by finding adjacent pairs of transcripts. A) RNA is present in a biological sample of interest. B) RNA molecules are barcoded and amplified locally, forming polymerase colonies (polonies). C) Where polonies overlap, their amplicons can be engineered to fuse together, forming concatemers. D) Sequencing concatemers reveals the adjacency of all polonies, which can be displayed in a graph. The number of concatemers formed between two polonies is the edge weight. From this graph, the relative spatial coordinates of each molecule can then be obtained.

However, experimental conditions can give rise to erroneous signals that create artifacts in the adjacency graph and disrupt the spatial reconstruction. We consider two such errors. The first of these is that of spurious crosslinks, formed between any pair of nodes regardless of position. These can be formed by incomplete PCR during post-experimental library preparation when products are no longer spatially confined, in a reaction similar to barcode-swapping and index-hopping events^11^. The second type of error is a fused node. When two polonies contain the same barcode or very similar barcodes that are misstakenly fused by sequencing error correction, they are represented in the adjacency graph as a single node, which can lead to distortions in the reconstruction.

Here, we propose two new methods to remove these errors, collectively called MinIPath (Minimum Indirect Path analysis). These methods are based on graph analysis on the adjacency graph alone. Spurious crosslinks are detected by finding a short indirect path connecting two directly connected nodes, which we show is harder to find if two nodes are far away and erroneously connected. Fused nodes are detected by looking at the connected nodes of any node. When these can easily be separated into two indirectly connected groups, the node is likely a fused node and can be split. We show the effect of both types of errors on the reconstruction quality using simulated diffusion-based data as input, and that these can be corrected by our method. In addition, we analyze a previously described DNA microscopy dataset^7^ and show that we can obtain accurate reconstructions by removing spurious crosslinks more efficiently than with a read count filter. In summary, this method provides a new, efficient way to filter artifacts from adjacency-based data, which can improve the overall quality of the resulting spatial reconstruction.

## Results

### A correction algorithm based on indirect paths removes spurious crosslinks and splits fused graph nodes

The effect of spurious crosslinks and node fusions on spatial reconstructions can be illustrated using a simple 2D grid as input (Fig. 2A), where each node is connected to their neighbors and each edge has a weight of 1. In line with previous work, we focused on undirected, bipartite, weighted graphs *G* for the extend of this study,

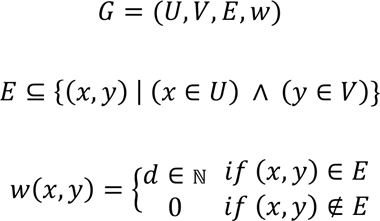

where *U* and *V* are the independent sets of nodes of the bipartite graph, *E* is the set of edges, and *w* is a mapping function assigning the edge weights. These graphs arise out of an experimental setup where two types of polonies are used that can only react with each other, to prevent the formation of concatemers within one polony^7^. Edge weights are equal to the number of concatemers formed between two polonies, found by barcoding each unique product formed. Spurious crosslinks connect any two nodes (Fig. 2B), while a fused node keeps all the edges of its original two nodes (Fig. 2C), regardless of the distance between them. Both types of errors greatly distort the resulting spatial reconstructions (Fig. 2D-F).

**Figure 2.**
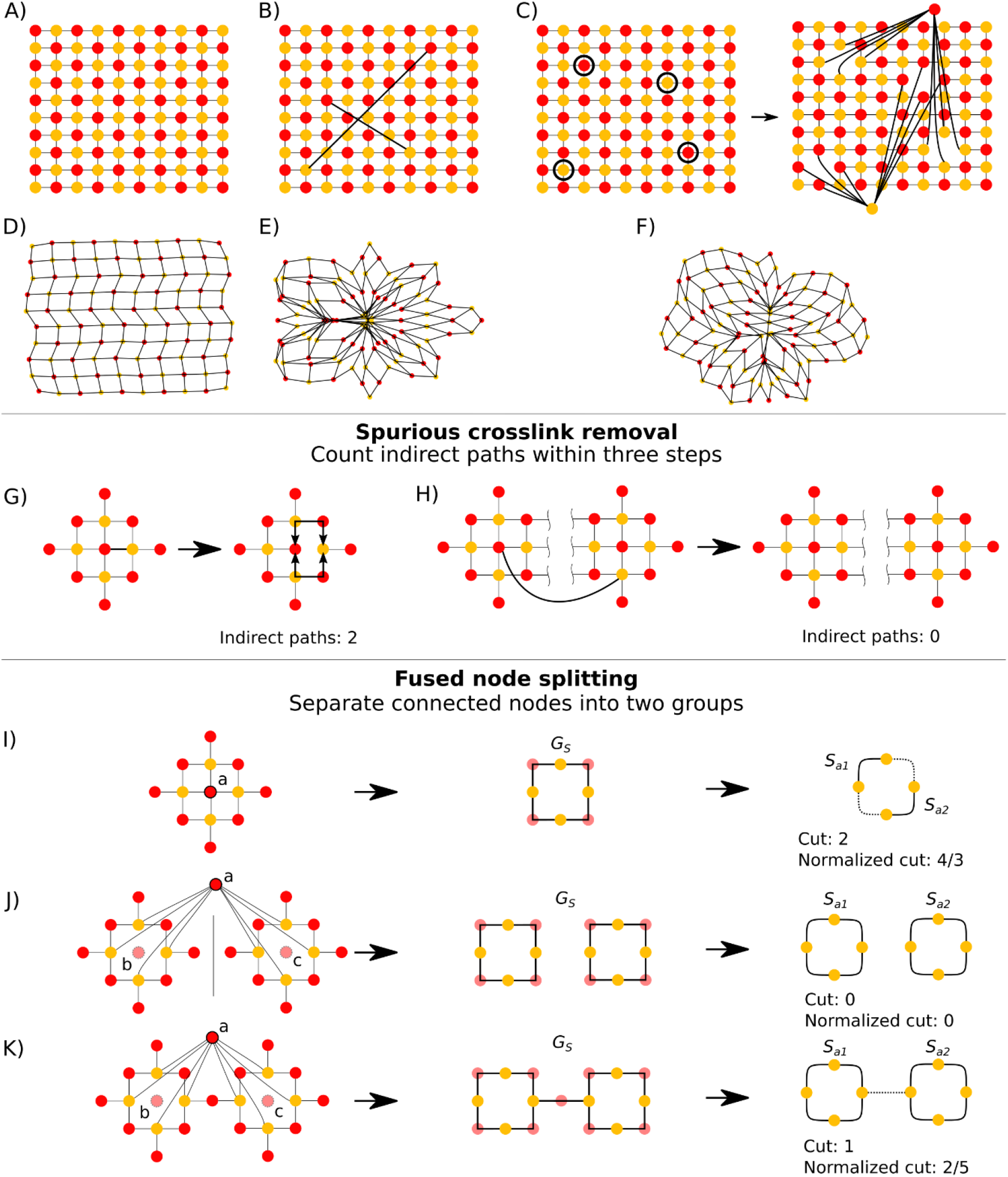
Spurious crosslinks and fused nodes can be corrected using methods based on indirect paths. A) Base grid used as an example. Each node is connected to their direct neighbours. B) as (A), but with two spurious crosslinks. C) as (A), but two sets of nodes, marked by circles, are fused. The two fused nodes maintain their connections. D-F) sMLE reconstruction of (A-C). G-H) Spurious crosslinks can be detected by counting the number of indirect paths at three steps. An edge connecting two neighbours has two indirect paths at three steps (G), while a spurious crosslink has no indirect path at three steps (H). I-K) Fused nodes can be detected by, for some node *a*, creating the graph *G_S_* out of the nodes that have edges to *a* and forming edges between if a path of two steps can be found between them in *G*, without using edges to *a*. If *a* is a non-fused node, partitioning *G_S_* requires the removal of 2/4 edges, resulting in a normalized cut of 4/3. If *a* is the result of fusing two nodes (*b* and *c*), partitioning *G_S_* requires the removal of fewer edges, and the partition has a lower normalized cut of 0 (J) or 2/5 (K).

Spurious crosslinks can be identified by counting the number of short indirect paths between two nodes connected by an edge. For an edge between two neighbours, one can find two indirect paths connecting them within three steps (the minimum in a bipartite graph; Fig. 2G). By contrast, for two nodes that share no direct neighbours, no indirect paths can be found within three steps (Fig. 2H). To remove spurious crosslinks, we therefore calculate the number of indirect paths connecting each edge in three steps.

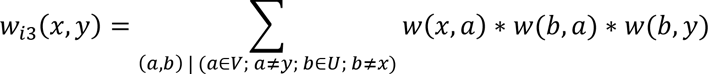

In unaltered graph (Fig. 2A), 𝑤_𝑖3_(𝑥, 𝑦) > 0 if *x* and *y* are neighbours. However, if x and y are not neighbours and are connected through a spurious crosslink, 𝑤_𝑖3_(𝑥, 𝑦) = 0. Calculating *w_i3_* therefore provides a metric to distinguish between spurious crosslinks and other edges.

For fused node correction, we further build on the intuition that spatially close nodes should be connected with short paths. Let *S_a_* denote the set of nodes with edges to some node *a*. Although these nodes do not have edges among themselves (since they belong to the same bipartition), they can be indirectly connected at two steps. For convenience, we form the graph *G_S_* from the nodes *S_a_*,

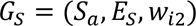

Where

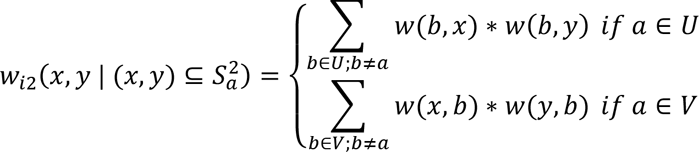

and *E_S_* is the set of edges between the nodes in *S_a_*. Note that the edges to the original node *a* are not used to find the edges in *w_i2_*.

If *a* is not a fused node (Fig. 2I), *G_S_* is connected. However, if *a* is a fused node, i.e., the result of fusing nodes *b* and *c*, then 𝑆_𝑎_ = 𝑆_𝑏_ ∪ 𝑆_𝑐_. There are no indirect edges between the nodes of *S_b_* and *S_c_* if *b* and *c* are sufficiently far apart (Fig. 2J). In the 2D example grid, if *G_S_* is not connected, we can therefore assume that node *a* is a fused node. To split it correctly, we simply remove node *a* and create two new nodes in *G* that each inherit the edges to the nodes in each of the connected groups in *G_S_*.

However, the method described above fails if even a single indirect edge is found between the nodes in *S_b_* and *S_c_*, e.g., if nodes *b* and *c* are within four steps of each other (Fig. 2K). To make our method more robust, we instead aim to partition *S_a_* into sets *S_a1_* and *S_a2_* so that *S_a1_* and *S_a2_* resemble *S_b_* and *S_c_* as closely as possible. Since we expect there to be more indirect edges within the nodes of *S_b_* and *S_c_* than between *S_b_* and *S_c_*, the problem can be thought of as obtaining a partition of *G_S_* that seeks to minimize the number of removed indirect edges between *S_a1_* and *S_a2_* while maximizing the number of remaining indirect edges within *S_a1_* and *S_a2_*. This is analogous to spectral graph partitioning^12^. To evaluate the partition, we calculate the normalized cut (ncut)^13^. The cut is defined as the sum of the edge weights removed between the *S_a1_* and *S_a2_* by the partitioning,

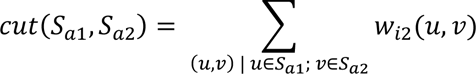

while the normalized cut equals the cut divided by the sum of the edge weights in each subgroup^13^.

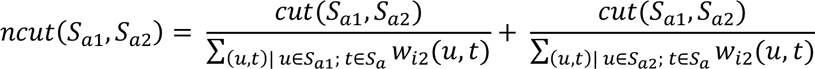

The normalized cut has a range of 0 to 2. When a single indirect edge is present between the connected groups (Fig. 2K), the partition of *S* has a low normalized cut (2/5). By contrast, if *a* is not a fused node (Fig. 2I), partitioning the graph *G_S_* requires the removal of more edges, resulting in a high normalized cut (4/3). The normalized cut therefore provides a continuous metric which can be used to distinguish fused from non-fused nodes, even when *G_S_* is connected.

### The error correction algorithm removes errors across a wide range of diffusion-based simulated adjacency graphs

To test our method on more complicated data, we used a model for polony formation in hydrogels to generate adjacency data. Starting from Fick’s law for diffusion for a singe polony, one can derive the relationship between the reaction rate *ω* between two polonies and their distance^7^:

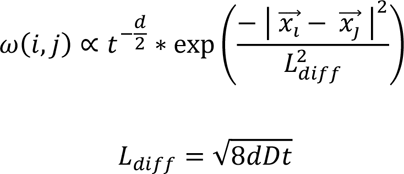

Where *ω(i, j)* is the reaction rate between two polonies *i* and *j*, *D* the diffusion constant, *t* the time since polony creation, *d* the number of dimensions, and *x_i_* and *x_j_* the location of polonies *i* and *j*, respectively. *L_diff_* describes the characteristic diffusion length and scale of the distribution. The exact proportionality constant is not known but depends on the inherent reactivity of two monomers.

As polony input, we created a 2D grid of nodes randomly distributed a shape (Fig. 3A; 200 x 450 pixels; ∼4000 nodes). Based on the diffusion model, the edge weight of any two nodes of different types was set to

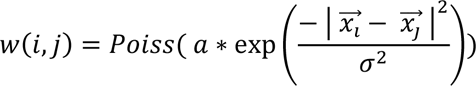

where *a* represents the amplitude and σ the spread. In experimental terms, the amplitude can be affected by the inherent reactivity of the polonies and sequencing depth, both of which determine how many products are seen. If the amplitude is high, multiple concatemers are formed and sequenced for each polony pair, resulting in larger edge weights in the resulting graph. The spread is analogous to *L_diff_* and determines the distance at which two polonies might still react. Experimentally, it will be determined primarily by the diffusion constant, and therefore by the properties of the hydrogel and size of the products (Fig. 3B).

**Figure 3.**
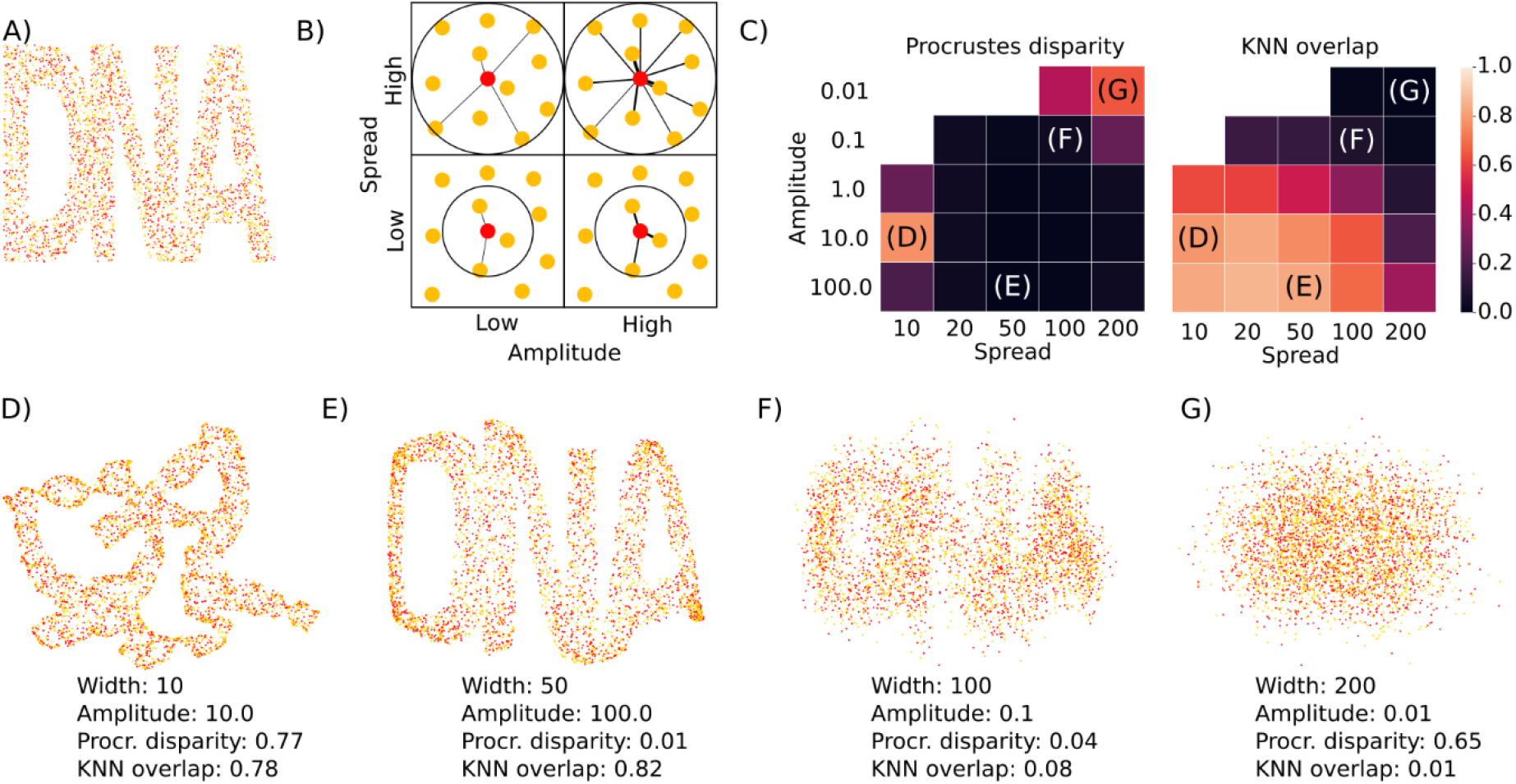
Reconstructions quality depends on Gaussian parameters that determine adjacency. (A) Node locations used as input, on a grid of 200 by 450 pixels. B) Edges are formed between neighbouring nodes. The spread determines the range at which edges are formed, while the amplitude determines the edge weight. C) The reconstruction quality depends on the amplitude and spread. Globally accurate reconstructions (low Procrustes disparity) are obtained unless the amplitude or spread are too low. Locally accuracy (high KNN overlap) increases with amplitude but decreases when spread approaches the sample size. (D – G): Example reconstructions with different spreads and amplitudes: (D) good local, poor global quality, (E) good local, good global quality, (F) poor local, good global quality and (G) poor local, poor global quality.

We generated adjacency data using a wide range of amplitudes and spreads and reconstructed the polony locations using the previously described spectral Maximum Likelihood Embedding (sMLE) method^7^. We reconstructed all adjacency datasets where at least 80% of all nodes were connected in a single group and evaluated all reconstructions using the Procrustes disparity as a global metric, and the number of overlapping neighbors out of the 15 nearest as a local metric^9^.

The different parameters used in the simulation had great influence on the reconstruction quality. Local accuracy depended primarily on the spread: if it was small (σ ≤ 50), local accuracy was high even when global accuracy was low (KNN ≥ 0.75; Fig. 3D), but when it started to approach the sample size, local accuracy decreased (Fig. 3F, 3G). Higher amplitudes also led to better local reconstructions, and the combination of low spread and high amplitude led to the most accurate reconstructions both globally and locally (Fig. 3E).

To examine the effect of errors on the reconstruction, spurious crosslinks were added (1-20% of total edge weights) and nodes were fused (1-20% of all node pairs). As before, adding in these errors distorted the reconstructions, affecting the reconstruction (e.g., Fig. 4A). The global reconstruction quality was affected more than the local reconstruction quality, as a small number of errors can twist the reconstruction without affecting the nearby neighbors of each point. Increasing the spread of in the simulation made the reconstructions more robust, but changing the amplitude had little effect on the robustness to errors (Ext. Data. 1).

**Figure 4.**
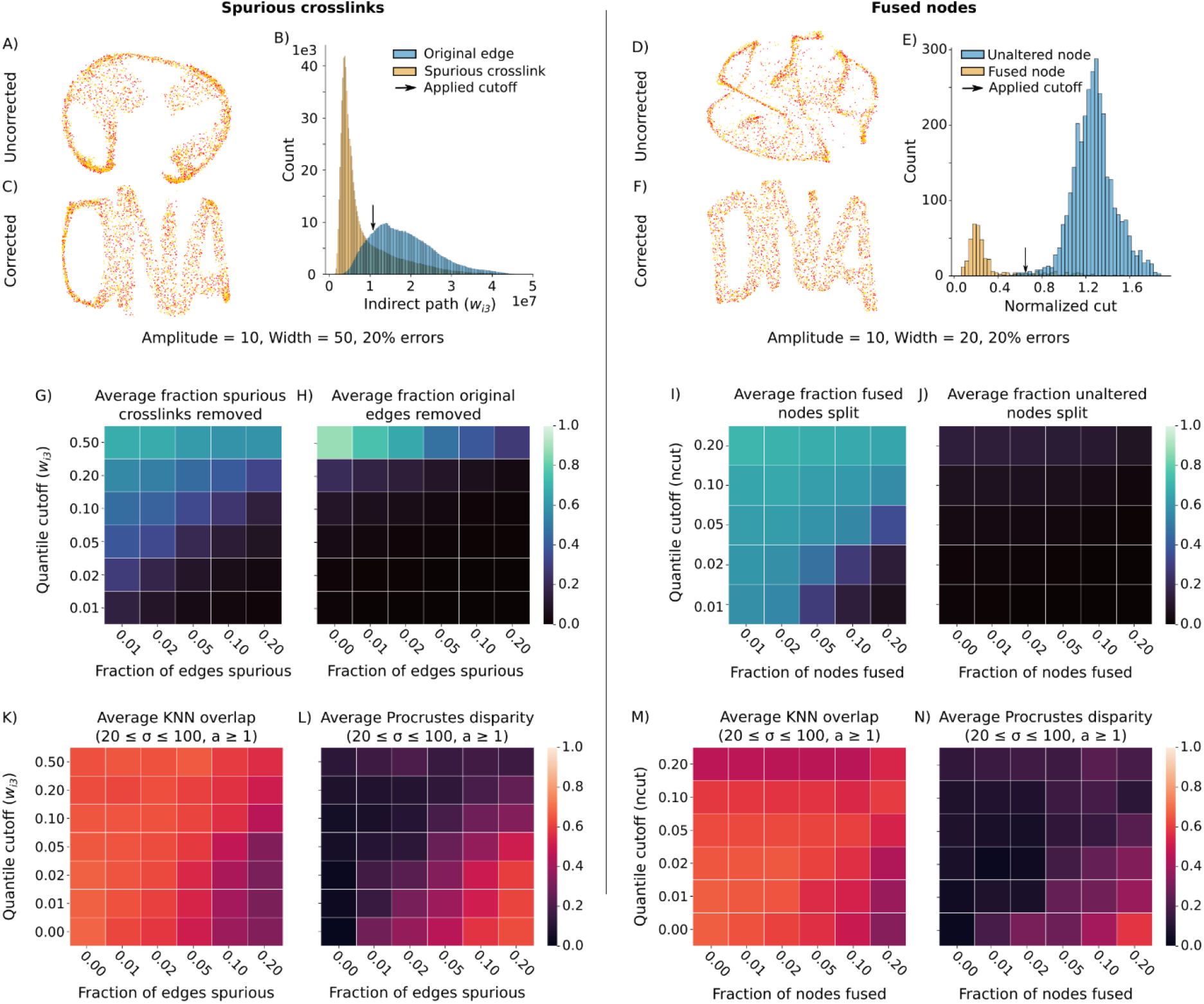
Indirect path analysis rescues reconstructions by removing spurious crosslinks and corrects fused nodes. (A, D) Example distorted reconstructions at amplitude 10, and width 50 and 20, respectively, with 20% errors. Arrows indicate condensed lines of points. (B, E) Indirect paths and normalized cuts distributions of the pairing data used to generate the reconstructions in A and D. The arrow indicates the cutoff used. (C, F) Reconstructions after correction. (G-J) Average fraction of true positive (G, I) and false positive (H, J) across all simulated datasets. (K-N) Average reconstruction qualities before and after corrections across simulated datasets that were affected by the errors (with amplitude of 1, 10 or 100 and width of 20, 50 or 100).

As before, we counted the indirect paths at three steps to identify spurious crosslinks. In contrast to the previous example, the number of indirect paths was not always 0 for spurious crosslinks, but instead depended on the amplitude, spread, number of other errors and distance between the nodes (Ext. Data. 2). However, since a spurious crosslink connects two randomly selected nodes, the distance between these nodes is larger on average than the distance between two nodes connected by a non-spurious edge. As a result, most spurious edges had lower indirect path values than other edges (Fig. 4B), meaning they could be identified, removed and the reconstruction restored (Fig. 4C).

Similarly, we used the fused node correction algorithm to detect and split fused nodes. Fused nodes could disrupt the reconstruction (example Fig. 4D). Identification of split nodes by forming *G_S_* for each node, partitioning it with spectral graph partitioning and evaluating the partition by calculating the normalized cut proved effective (Fig. 4E). The wider spread of connections meant that *G_S_* was not disconnected for most nodes, but their partitions had lower normalized cut values than those of unaltered nodes. Splitting up nodes below a certain cutoff again restored the reconstruction (Fig. 4F).

On average across all simulations, the indirect-path-based correction algorithm preferentially corrected the spurious crosslinks and nodes over original edges and unaltered nodes (Fig. 4G-4I). The ratio of true positive to false positive depended primarily on the spread, i.e., spurious crosslinks and fused nodes were more easily identified when only local reactions were formed (Ext. Data. 3). However, applying the algorithm without the presence of errors did not affect the reconstructions, except when an exceptionally high cutoff (quantile of 0.50) was used, suggesting that the error correction algorithm can safely be used on unaltered data without affecting reconstruction quality.

Although not all reconstructions were affected by the errors, all reconstructions that were affected could be improved with the correction algorithm (Ext. Data. 4-7). Zooming in on simulations that were initially accurate but strongly affected by the errors (20 ≤ spread ≤ 100, amplitude ≥ 1.0), the average Procrustes disparity increased from 0.02 +- 0.03 to 0.63 +- 0.10 and to 0.52 +- 0.12, when spurious crosslinks or fused nodes were added, respectively. However, the average quality improved by applying the correction algorithm (Fig. 4K-4N), and for each case where the introduced errors affected the reconstruction, a cutoff could be found that restored it (Ext. Data. 4-7). When fused nodes were corrected, we evaluated the node splitting accuracy, i.e., whether the nodes after splitting were connected to the same nodes as before the nodes were fused (see Methods). We found that the resulting groups matched accurately to the original (average Jaccard index: 0.73 +- 0.17), and most accurately when the spread in the simulation was low (Ext. Data. 8). Node splitting only proved ineffective when spread and amplitude were low. In these cases, *G_S_* consisted of more than two components, and the node split could not be evaluated (Ext. Data. 8).

In summary, the proposed method removes spurious crosslinks and corrects fused nodes across a wide range of simulated data, even in the presence of up to 20% spurious crosslinks of 20% fused nodes.

### Applying the correction algorithm on experimental data efficiently removes disruptive crosslinks

To see how our method would perform on experimental data, we analyzed a previously published DNA microscopy dataset for which a reference image was available^7^. In this experimental setup, specific types of RNA transcripts (*actb* for “beacons”, and *gfp, rfp,* and *gapdh* as “targets”) were used as seeds for the two types of polonies in the bipartite graph. Each product connecting two polonies could be recognized by a unique barcode called a Unique Event Identifier (UEI), the number of which was used as edge weight.

Using the same pipeline as described before, we extracted 1.26E5 polonies with 6.72E5 edges and 9.55E5 unique UEIs from the raw sequencing data. When using all data to reconstruct the largest connected group, the resulting reconstructions frequently collapsed into a “star-like” pattern (Ext. Data. 9). Only one out of ten reconstructions produced a layout that could be overlayed on the microscopy image with a poor match (Fig. 5B). To remove possible artifacts, Weinstein *et al* applied a read count filter that removed all products without sufficient reads. While this strategy did improve the reconstruction quality, a read count filter of 4 was required to obtain an accurate reconstruction (Fig. 5C and 5D). Given the large number of products produced during a DNA microscopy reaction, many of them had low read counts (Ext. Data. 10), and as a result, only 70.6% (6.39E5/9.03E5) of all UEIs remained for reconstruction.

**Figure 5.**
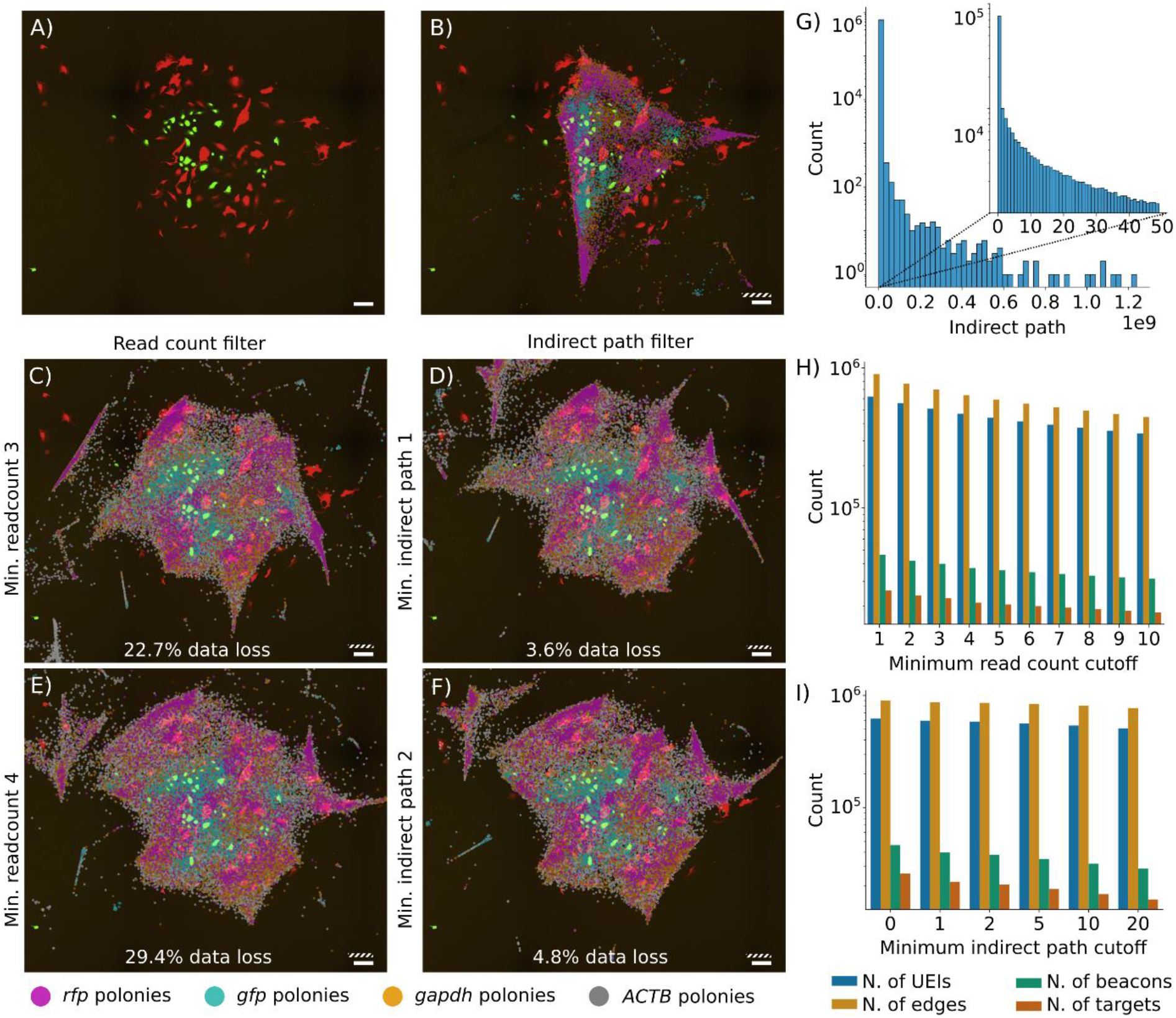
Applying a minimum indirect path filter efficiently removes topological artifacts from experimental data. Data loss is calculated as the total number of UEIs remaining for reconstruction after the applied filter. A) Reference microscopy image, adapted from^7^. B) Reconstruction from uncorrected and unfiltered data. C, E) Reconstructions after read count filters of 3 and 4 respectively. D, F) Reconstructions after indirect path filters of 1 and 2 respectively. G) Indirect path distribution. H, I) Number of remaining UEIs, edges, and beacon and target polonies after the read count filter and the indirect path filter, respectively.

By contrast, applying a minimum indirect path cutoff of 1 greatly improved the reconstruction quality, while only removing 3.6% of all UEIs (3.2E4/9.03E5; Fig. 5E). The reconstruction quality further improved with an indirect path cutoff of 2 (removing 4.8% of all UEIs; Fig. 5F). Applying an even higher cutoff did not clearly further improve the reconstruction. Using an indirect path cutoff therefore removed disruptive edges more efficiently than a read count filter, allowing more data to be used for the resulting reconstruction.

We additionally attempted to split possible fused nodes in this dataset. Like simulated data with low amplitude and spread, *G_S_* was disconnected into more than two components for most of the nodes (8.9E4/1.1E5; 82.4%), meaning it could not be used for accurate partitioning. Indeed, when describing the experimental data in terms of amplitude and spread, we found it had low amplitude and average spread. The dimensions of the sample of an accurate reconstruction (Fig. 5F) were approximately 9x8 *L_diff_*^2^ equivalents, i.e., 11-13 times as long as the average distance of two connected nodes (0.68 +- 0.59 *L_diff_*^2^ equivalents), suggesting an average to low spread. Of all 1.94E7 polony pairs that were within the average pairing distance, only 6.72E5 edges (3.5%) were obtained from the sequencing data, similar to a low amplitude in the simulation. Possible fused nodes could therefore not be identified.

Overall, applying a minimum indirect path filter efficiently removed disruptive crosslinks from the experimental data, losing only a small percentage of all connections. Since the fidelity of the reconstruction scales with the amount of available data, the proposed algorithm could provide a useful filter to obtain more accurate reconstructions from sparse data.

## Discussion

DNA microscopy has the potential to accelerate biological research by providing new methods to investigate the spatial organization of transcripts and other molecules. Since the input data is paired sequencing data, this brings about new types of errors and noise that are specific to this method and for which no filtering or correction methods have been described so far.

In this work, we describe a method aimed at bridging this gap. The error correction algorithm presented here corrects two types of errors: spurious crosslinks and fused nodes. Spurious crosslinks are distinguished from real edges by counting the number of indirect paths at three steps, while nodes are split if they are connected to nodes in different topological environments, using spectral graph partitioning and the normalized cut as a continuous metric to identify them. Both methods work well on simulated data, even in the presence of up to 20% errors: most of the introduced errors could be identified and corrected, restoring the spatial reconstructions.

Although we limited our study to indirect paths at three steps for spurious crosslink removal, it is possible that longer indirect paths (five, seven or more steps) could provide a more accurate distinction between spurious crosslinks and other edges. However, this comes at a higher computational cost. Calculating the indirect paths at three steps has a worst-case performance of 𝑂(|𝐸|^3^ ∗ |𝑈 ∪ 𝑉|), which increases to 𝑂(|𝐸|^5^ ∗ |𝑈 ∪ 𝑉|) or 𝑂(|𝐸|^7^ ∗ |𝑈 ∪ 𝑉|) for five or seven steps, respectively. Calculating these paths could therefore become prohibitive for large experimental datasets.

The experimental data could not be directly reconstructed, but required filtering of pairs before an accurate reconstruction could be obtained. These results highly suggest that a significant fraction of the paired data are indeed experimental artifacts that disrupt the reconstruction. Although they could also be removed with a read count filter, applying the spurious crosslink removal algorithm proved a more efficient way to remove these artifacts. Applying a minimum indirect path filter could therefore become a valuable addition to DNA microscopy pipelines. Unfortunately, the experimental dataset proved too sparse to attempt node splitting. However, the simulated data shows that reconstructions can be robust to these errors up to a few percent. Given the barcode length of the experiment, these errors are expected to be present under this threshold and not significantly disrupt the reconstructions.

Although we have applied our error correction algorithm here on DNA microscopy data, we note that the same principle could be applied onto any dataset where adjacency is the primary source of data, such as Hi-C data^14^. Several methods have been described to obtain the 3D organization from pairing data between genomic regions^15^, and it remains unclear, to our knowledge, how artifacts affect these. For this purpose, the method could be adapted to work on non-bipartite graphs. How errors affect these reconstructions and if this correction algorithm can improve them remains a topic for future studies.

In summary, we present an error correction algorithm for adjacency data, that can detect spurious crosslinks and fused nodes, and correct them, improving reconstruction quality in both simulated data, and proving to be an efficient error correction tool for experimental data.

## Methods and Protocols

### Algorithm implementation

Two methods were implemented in python to calculate indirect paths. The first method starts from the asymmetric adjacency matrix 𝐴(𝑖, 𝑗) = 𝑤(𝑖, 𝑗), then calculating the three-step adjacency matrix 𝐴_3_(𝑖, 𝑗) = 𝐴 ∗ 𝐴^𝑇^ ∗ 𝐴, then, for each node pair with an edge, subtracting the paths that use their direct edge, while setting other node pairs to 0:

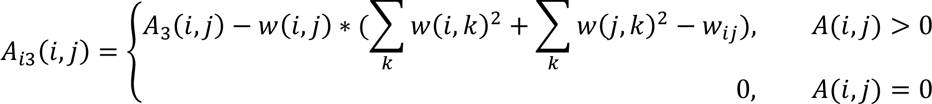

Calculating *A_3_* becomes computationally too complex for large experimental datasets. To this end, we implemented a version uses sparse matrices. It first calculates 𝐴_2_(𝑖, 𝑗) = 𝐴 ∗ 𝐴^𝑇^, which contains all two-step paths between all nodes of one partition (e.g., *U*). Then it iterates over every edge to find all indirect paths.

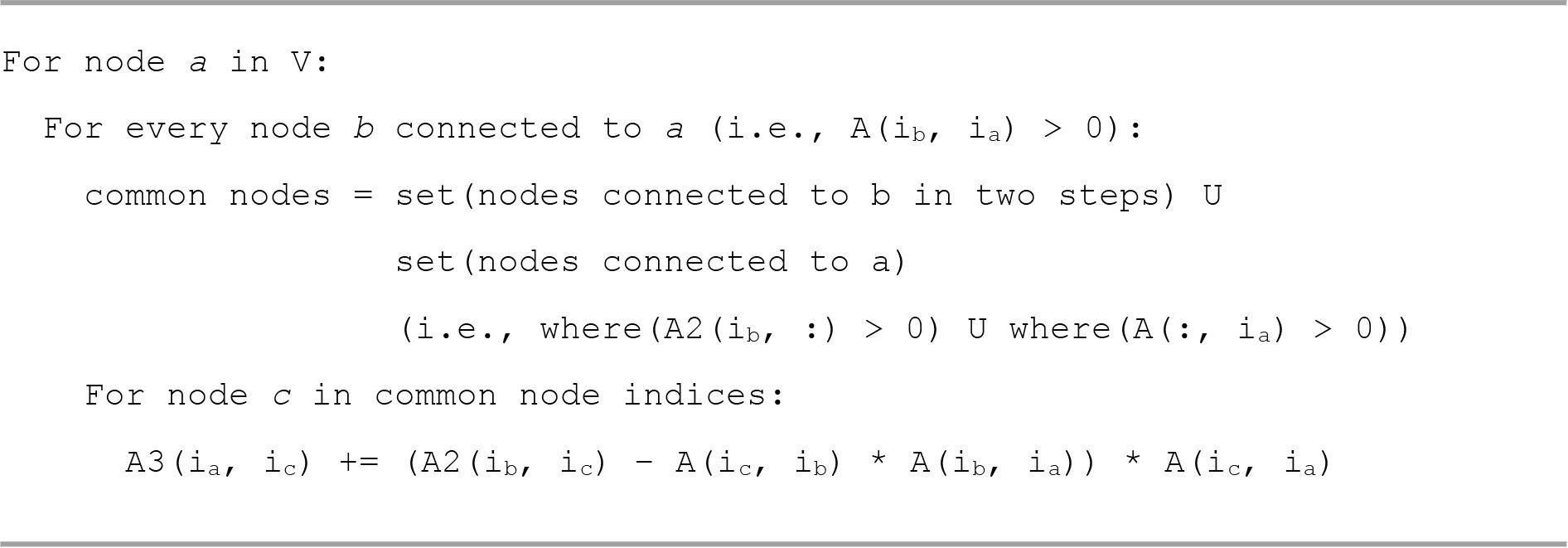

Where *i_x_* denotes the index of node *x*. The second method is implemented in python with Numba^16^ acceleration to allow for the use of multiple threads.

For node splitting, the graph *G_S_* was extracted and partitioned using spectral graph partitioning tool from the scikit-learn package^17^. Nodes were not considered for splitting if they had either fewer than four connections, or if *G_S_* consisted of more than two components. Normalized cuts were calculated first for all nodes, then nodes were selected for splitting according to the applied cutoff. If two nodes with an edge were both split, the edge was removed.

### Simulated data processing

Polony locations were reconstructed using the sMLE method described previously (original pipeline found at https://github.com/jaweinst/dnamic). A slightly adapted version of the pipeline was used to process large amounts of files more easily. In contrast to experimental data, simulated data was not subjected to the iterative minimum UEI filter prior to reconstruction. Reconstructions from graphs where the largest connected component was smaller than 80% of all nodes were not considered for further analysis. Global reconstruction quality was assessed using the Procrustes disparity, while local reconstruction quality was assessed by the overlap the k-nearest neighbours for each node in the original and reconstruction positions, as suggested earlier^9^.

To evaluate node splitting accuracy, we paired each set of nodes *S_a1_* and *S_a2_* to their closest match in *S_b_* and *S_c_*, and calculated the overlap:

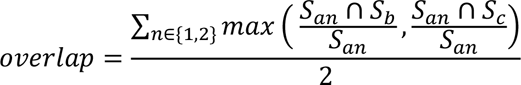

### Experimental data processing

Raw sequencing data was downloaded from the Sequencing Read Archive (project number PRJNA487001, sample 3), and processed as described previously, without a minimum read count. After applying a read count filter or indirect path filter, the remaining nodes were filtered as described earlier, first by iteratively removing nodes with less than two associated products (UEIs) to remove possible uncorrected sequencing errors, then by selecting the largest connected component. Data in Figure 5H-I describes properties of the graph after all filters were applied.

## Extended data

**Extended data 1.**
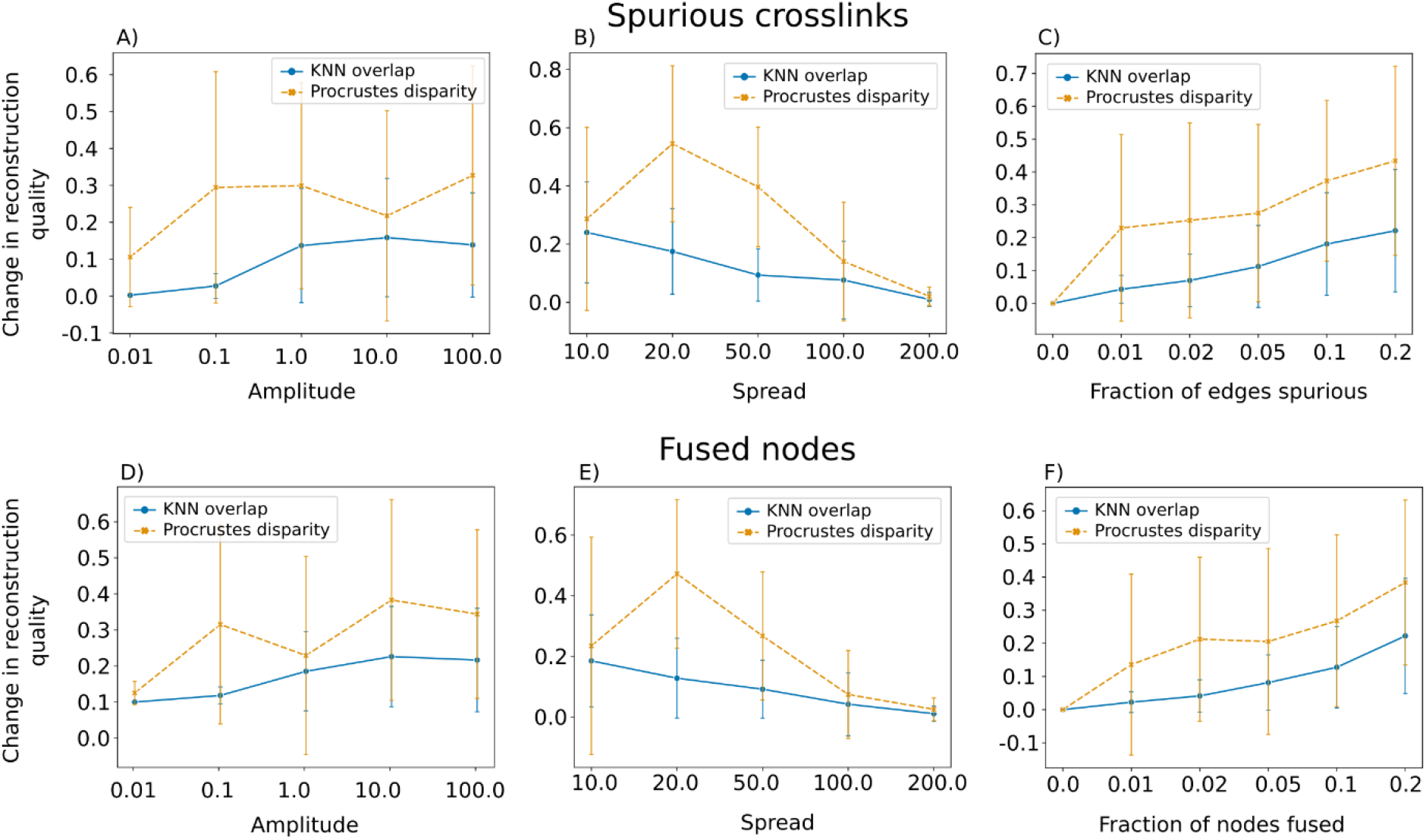
The robustness of reconstructions to introduced errors depends on the amplitude, spread and number of errors. Adding in either spurious crosslinks (A-C) or node fusions (D-F) decreases the reconstruction quality, on average. (A, D) At low amplitudes (≤ 0.1), reconstructions are generally of poor quality, which therefore does not change when errors are added. At high amplitudes (≥1.0), reconstructions were affected by introduced errors, but no clear change can be seen when increasing the amplitude from 1.0 to 100.0. (B, E) Increasing the spread allows for more connections to be formed, which makes the reconstruction more resilient to errors. (C, F) Increasing the number of errors also increases the severity of the disruption.

**Extended data 2.**
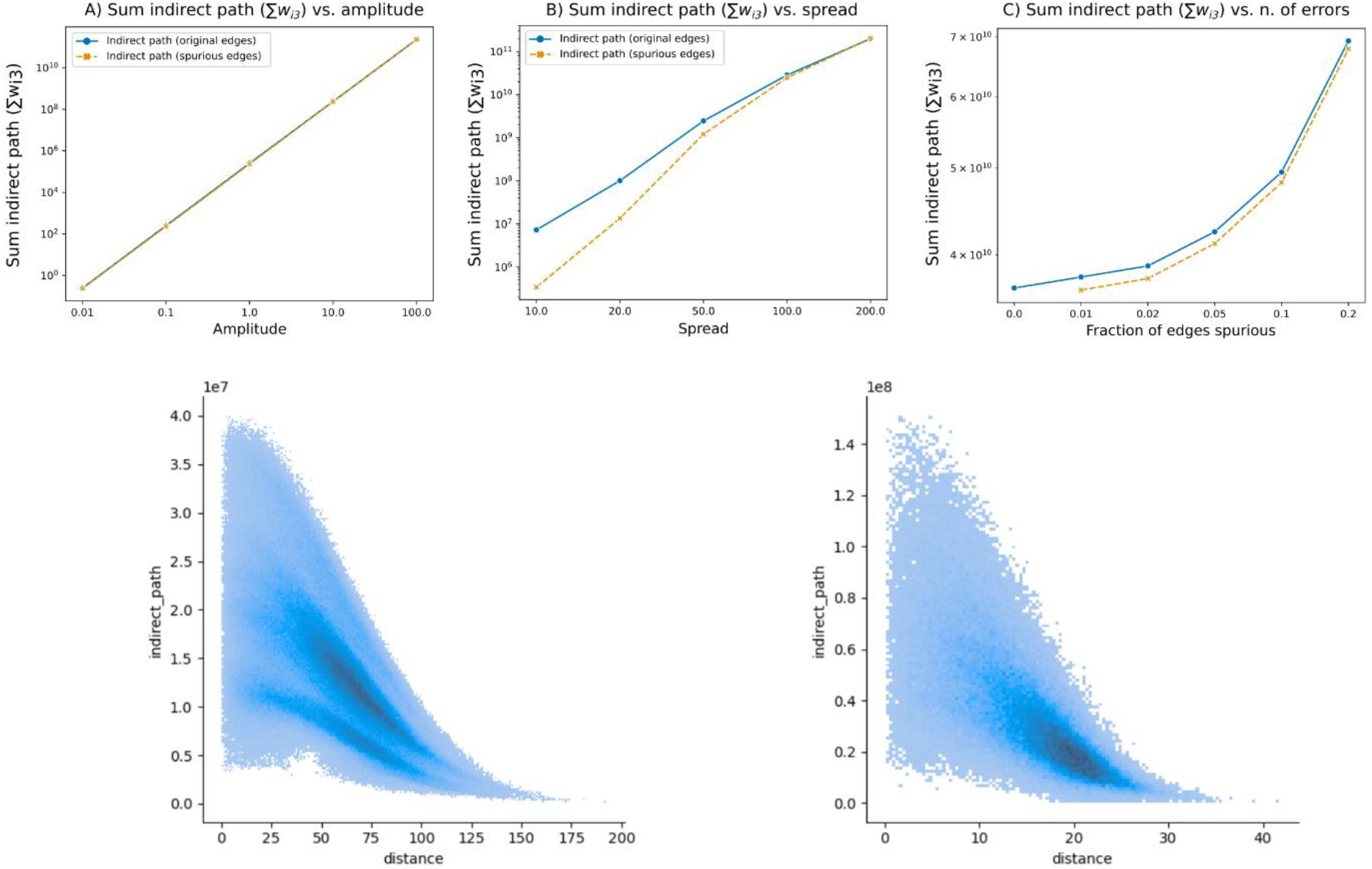
Indirect path increases with amplitude, spread, number of errors and distance. The total value of all indirect paths calculated across all simulations increases with A) amplitude, B) spread, and C) number of errors. Furthermore, the indirect path values at three steps decrease with distance between the polonies (D) Amplitude=10, spread=50. (E) Amplitude=10, spread=100.

**Extended data 3.**
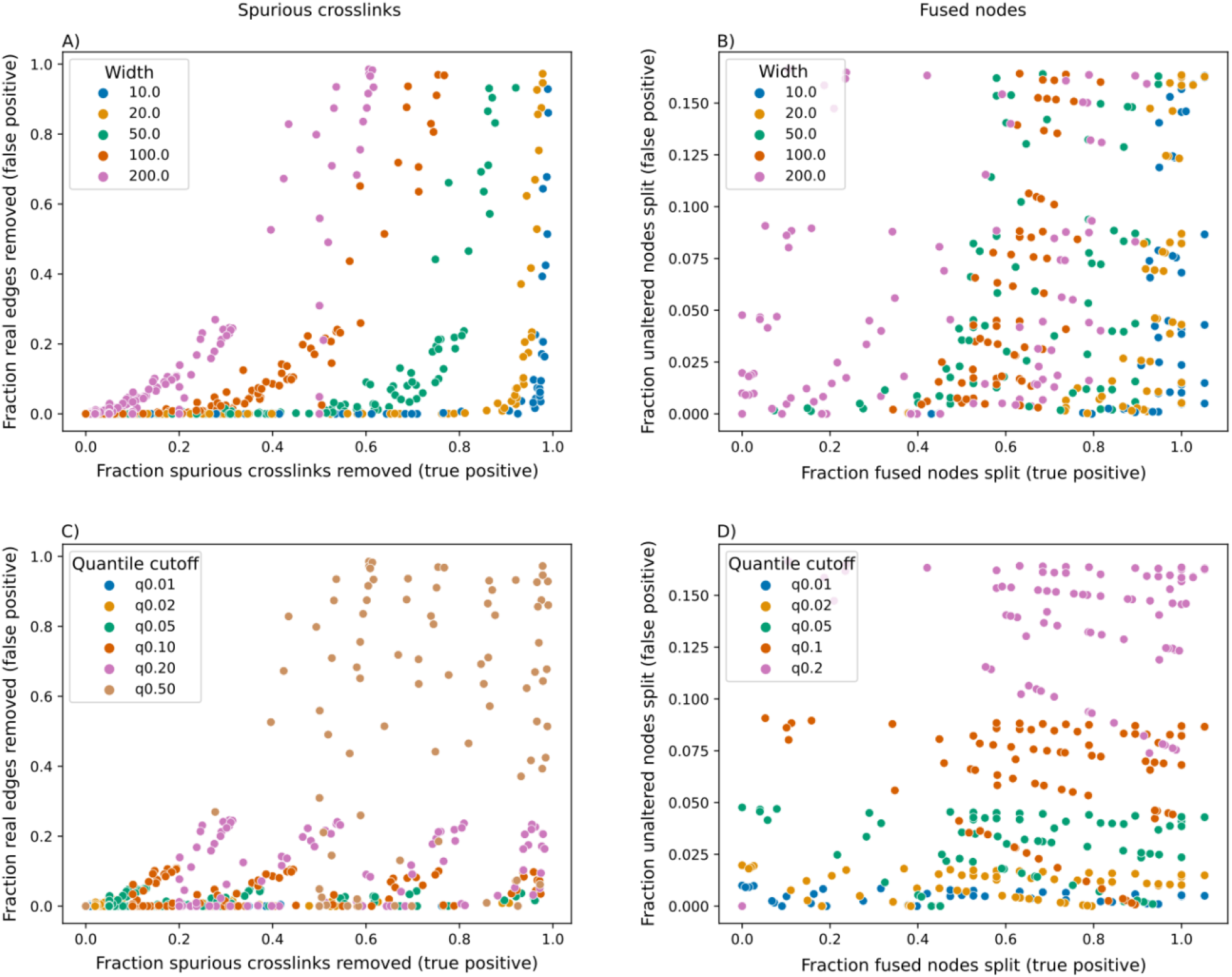
True positive versus false positive rate primarily depends on spread. Each point in each scatterplot represents the true positive and false positive rate in a single simulation where errors were added and corrected. (A, B) are colored by spread, (C, D) are colored by cutoff. The best ratio of true positive to false positive is reached with low quantile cutoffs and low spread.

**Extended data 4.**
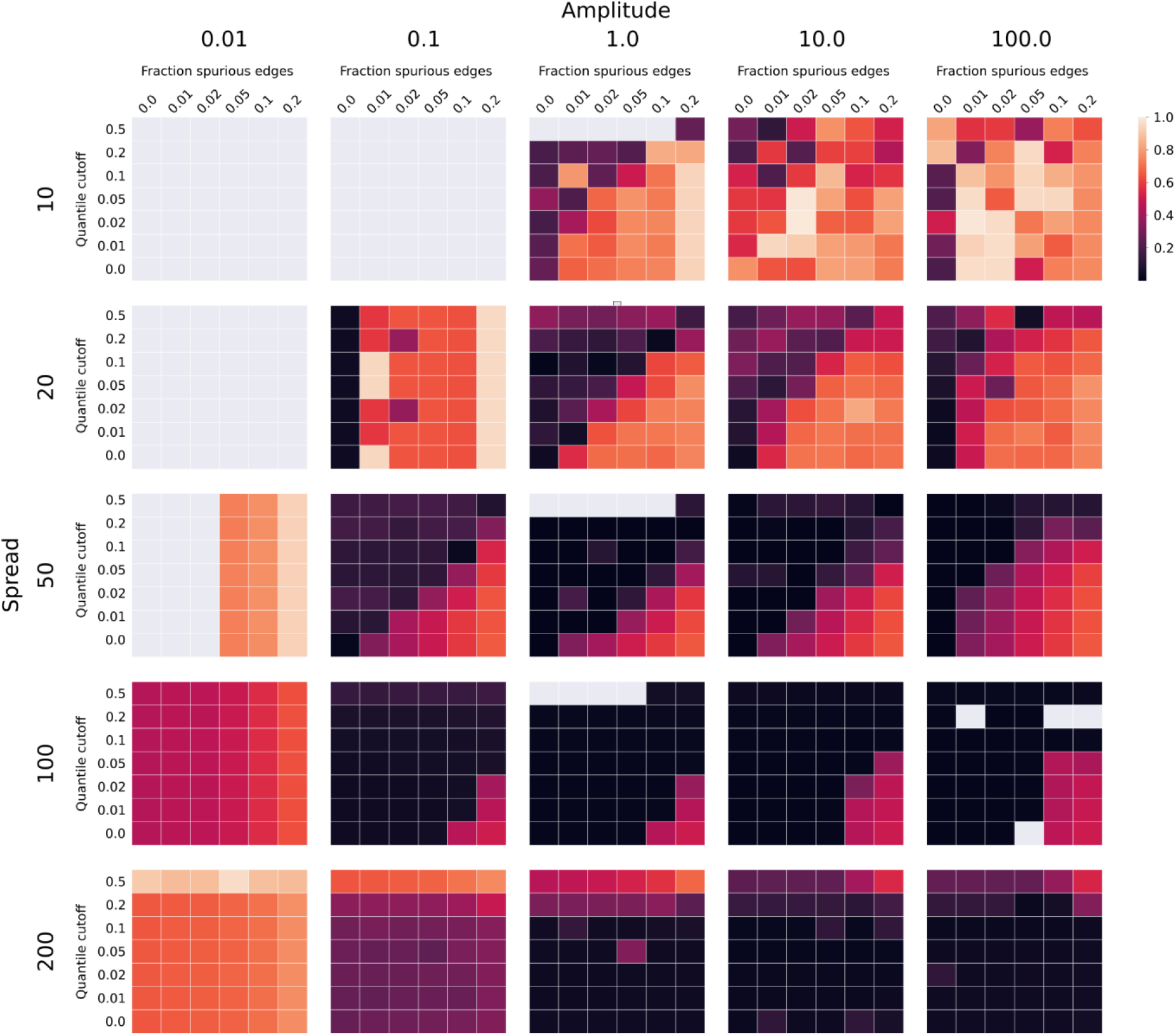
Procrustes disparity of reconstructions after spurious crosslinks are added and corrected across all simulations.

**Extended data 5.**
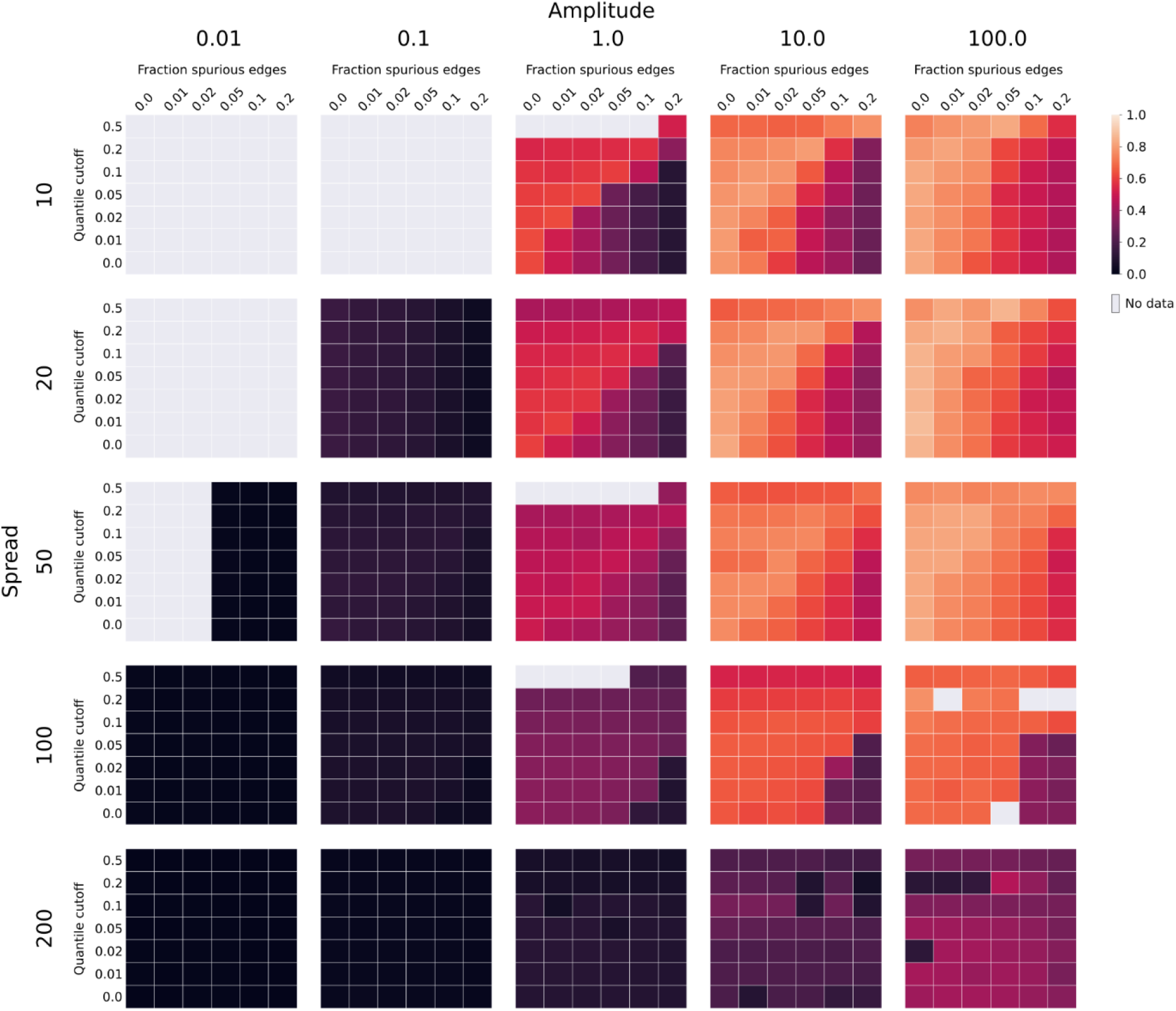
KNN overlap of reconstructions after spurious crosslinks are added and corrected across all simulations.

**Extended data 6.**
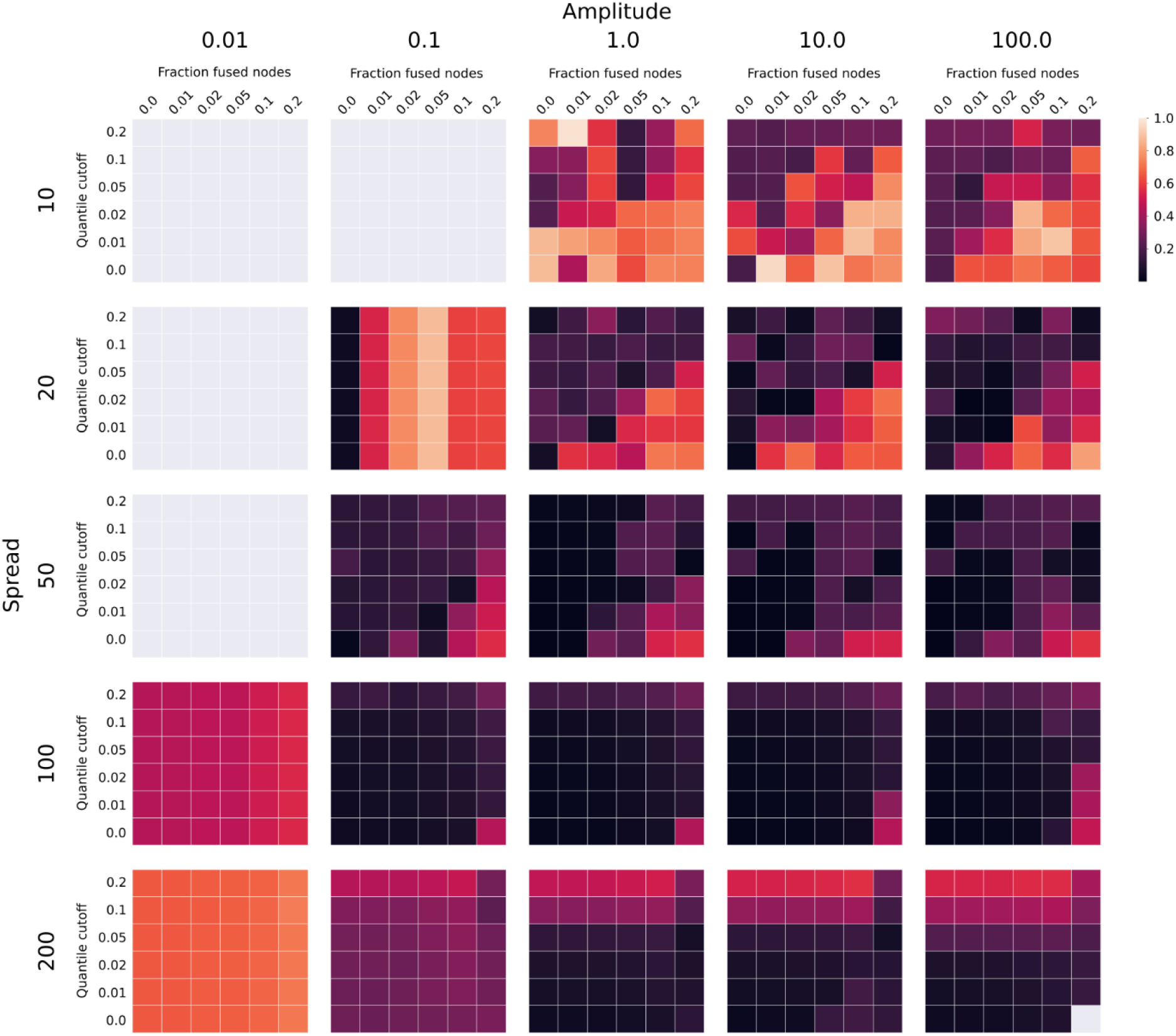
Procrustes disparity of reconstructions after nodes are fused and corrected across all simulations.

**Extended data 7.**
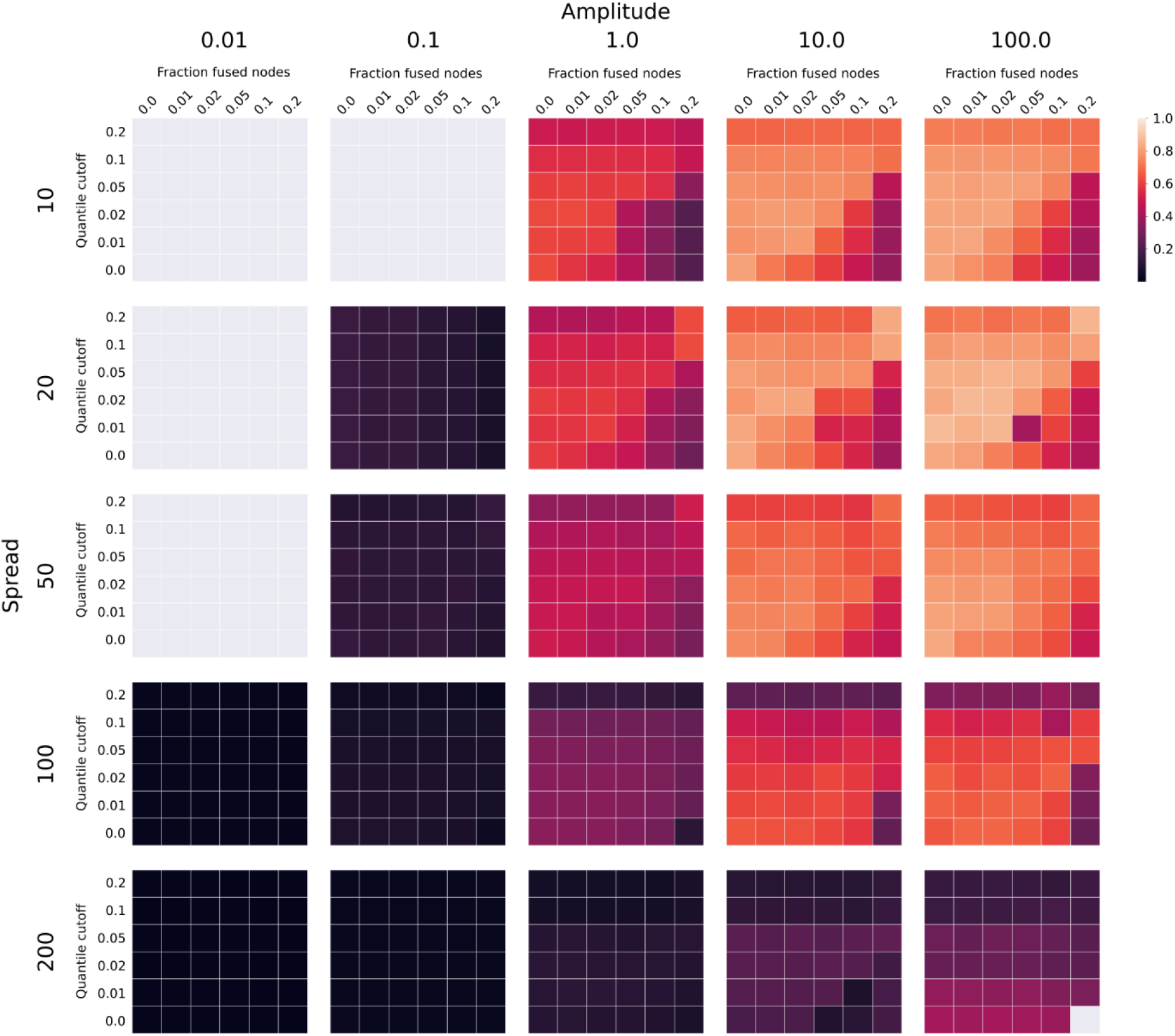
KNN overlap of reconstructions after nodes are fused and corrected across all simulations.

**Extended data 8.**
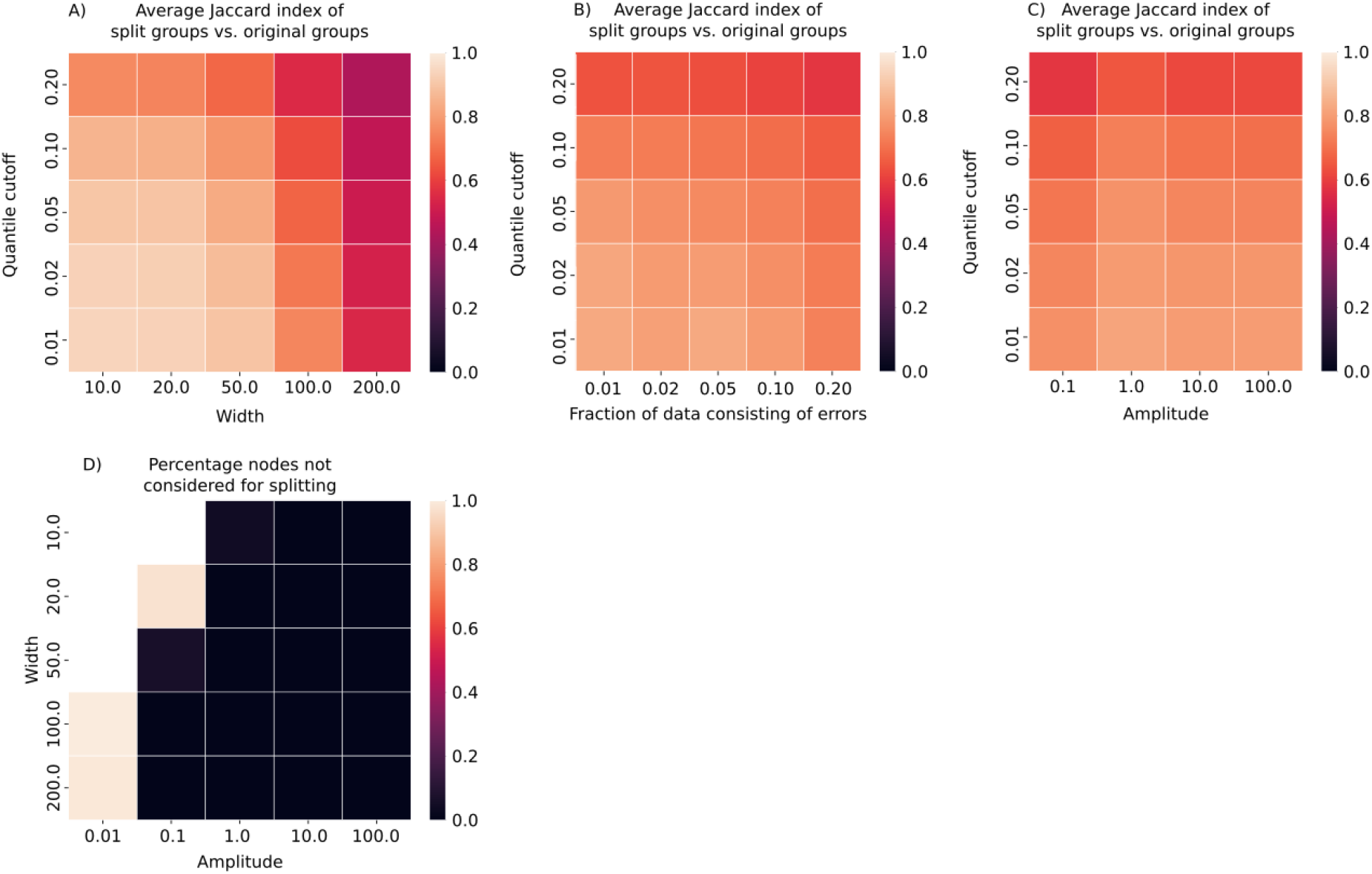
Further details on node splitting accuracy. The accuracy of node splitting was determined by matching each of the two partitions of connected nodes (*S_a1_* and *S_a2_*) after splitting to the original two groups of nodes connected to the nodes that were fused together (Sb and Sc), calculating the overlap of the best matching group, and averaging them. The overlap was the highest when the spread was low (A), became slightly less accurate as more errors were added (B), but did not depend on the amplitude in the simulation (C). (D) Nodes were only considered for splitting if they had at least four edges, and their connected nodes (*G_S_*) did not form more than two indirectly connected components. Generating data using low amplitude resulted in sparse data, where most of the nodes failed this criterium.

**Extended data 9.**
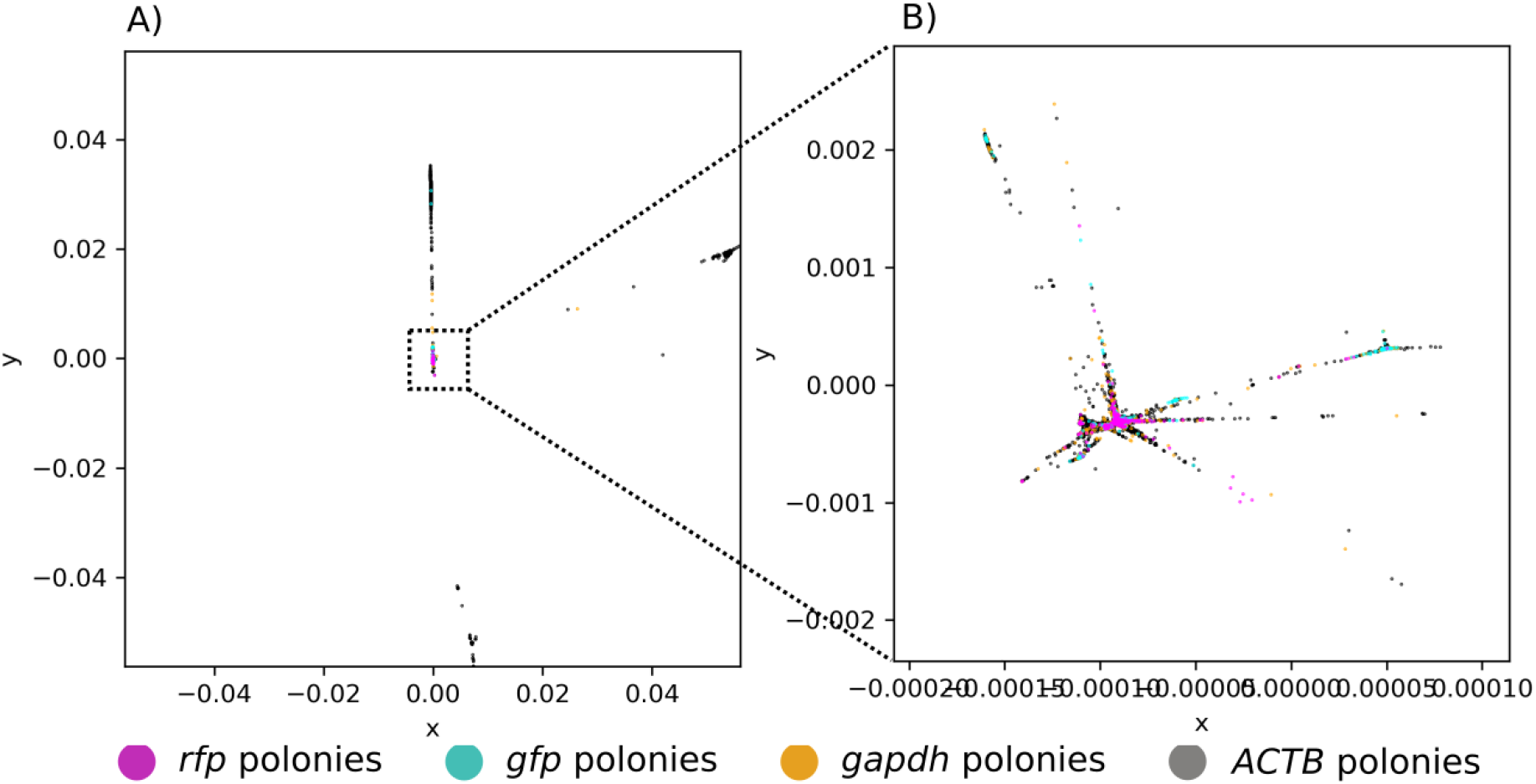
Uncorrected experimental datasets frequently collapsed during reconstruction. When unfiltered experimental data was reconstructed, the resulting reconstruction was frequently arranged in a star-like pattern. A) Full scale. B) Zoomed in.

**Extended data 10.**
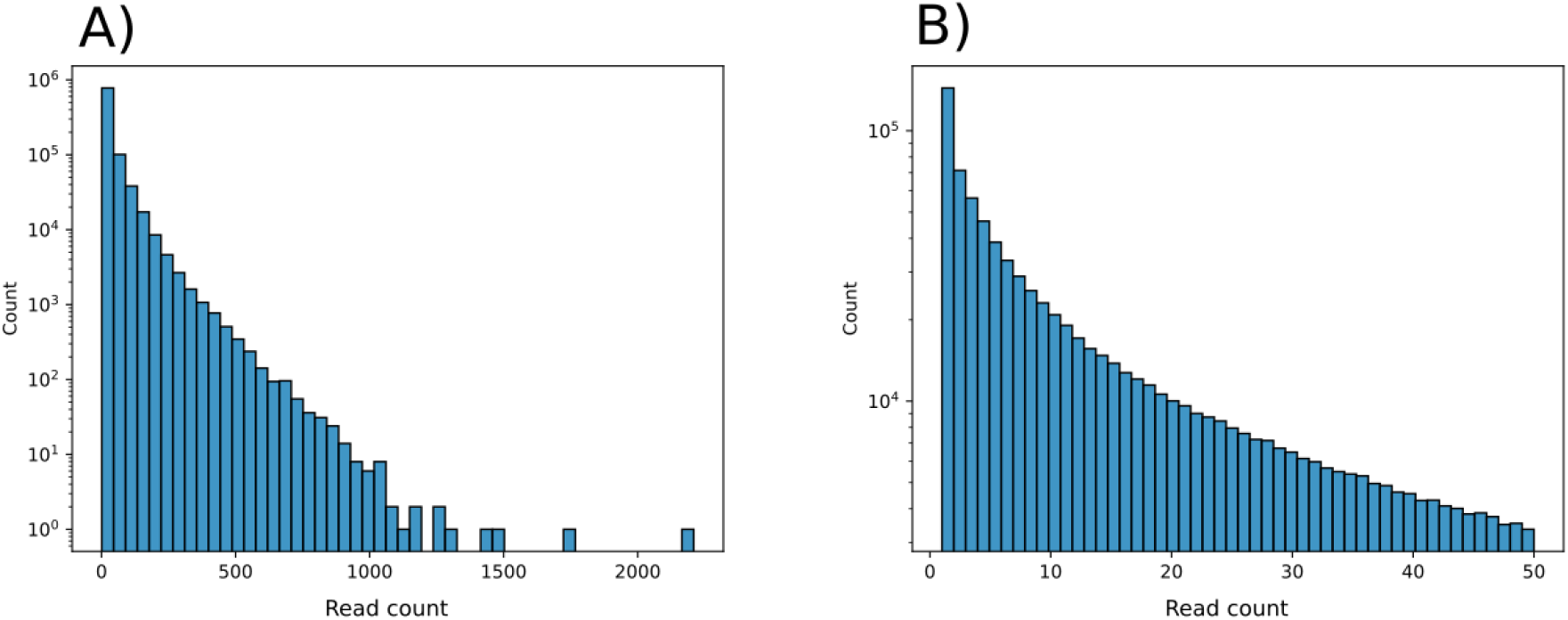
Read count distribution of the analysed experimental dataset. Although products were detected with read counts of more than 2000, the most abundant read count was 1. A) Full scale. B) Zoomed in on the range 1-50.

## References

1. Paolillo C, Londin E, Fortina P. Single-Cell Genomics. Clin Chem. 2019;65(8):972–985. doi:10.1373/clinchem.2017.283895

2. Moor AE, Itzkovitz S. Spatial transcriptomics: paving the way for tissue-level systems biology. Curr Opin Biotechnol. 2017;46:126–133. doi:https://doi.org/10.1016/j.copbio.2017.02.004

3. Moses L, Pachter L. Museum of spatial transcriptomics. Nat Methods. 2022;19(5):534–546. doi:10.1038/s41592-022-01409-2

4. Levy-Jurgenson A, Tekpli X, Kristensen VN, Yakhini Z. Spatial transcriptomics inferred from pathology whole-slide images links tumor heterogeneity to survival in breast and lung cancer. Sci Rep. 2020;10(1):18802. doi:10.1038/s41598-020-75708-z

5. Yoosuf N, Navarro JF, Salmén F, Ståhl PL, Daub CO. Identification and transfer of spatial transcriptomics signatures for cancer diagnosis. Breast Cancer Res. 2020;22(1):6. doi:10.1186/s13058-019-1242-9

6. Hoffecker IT, Yang Y, Bernardinelli G, Orponen P, Högberg B. A computational framework for DNA sequencing microscopy. Proceedings of the National Academy of Sciences. 2019;116(39):19282 LP - 19287. doi:10.1073/pnas.1821178116

7. Weinstein JA, Regev A, Zhang F. DNA Microscopy: Optics-free Spatio-genetic Imaging by a Stand-Alone Chemical Reaction. Cell. 2019;178(1):229–241.e16. doi:https://doi.org/10.1016/j.cell.2019.05.019

8. Boulgakov AA, Xiong E, Bhadra S, Ellington AD, Marcotte EM. From Space to Sequence and Back Again: Iterative DNA Proximity Ligation and its Applications to DNA-Based Imaging. *bioRxiv*. Published online January 1, 2018:470211. doi:10.1101/470211

9. Fernandez Bonet D, Hoffecker IT. Image recovery from unknown network mechanisms for DNA sequencing-based microscopy. Nanoscale. 2023;15(18):8153–8157. doi:10.1039/D2NR05435C

10. Mitra RD, Church GM. In situ localized amplification and contact replication of many individual DNA molecules. Nucleic Acids Res. 1999;27(24):e34–e39. doi:10.1093/nar/27.24.e34

11. Griffiths JA, Richard AC, Bach K, Lun ATL, Marioni JC. Detection and removal of barcode swapping in single-cell RNA-seq data. Nat Commun. 2018;9(1):2667. doi:10.1038/s41467-018-05083-x

12. von Luxburg U. A tutorial on spectral clustering. Stat Comput. 2007;17(4):395–416. doi:10.1007/s11222-007-9033-z

13. Shi J, Malik J. Normalized cuts and image segmentation. IEEE Trans Pattern Anal Mach Intell. 2000;22(8):888–905. doi:10.1109/34.868688

14. van Berkum NL, Lieberman-Aiden E, Williams L, et al. Hi-C: a method to study the three-dimensional architecture of genomes. J Vis Exp. 2010;(39). doi:10.3791/1869

15. Oluwadare O, Highsmith M, Cheng J. An Overview of Methods for Reconstructing 3-D Chromosome and Genome Structures from Hi-C Data. Biol Proced Online. 2019;21(1):7. doi:10.1186/s12575-019-0094-0

16. Lam SK, Pitrou A, Seibert S. Numba: A llvm-based python jit compiler. In: Proceedings of the Second Workshop on the LLVM Compiler Infrastructure in HPC.; 2015:1–6.

17. Pedregosa F, Varoquaux G, Gramfort A, et al. Scikit-learn: Machine Learning in {P}ython. Journal of Machine Learning Research. 2011;12:2825–2830.

